# Chain length defines the spatial segregation and metabolic fate of fatty acids in adipocytes

**DOI:** 10.1101/2025.10.28.685119

**Authors:** Dávid Kovács, Romain Gautier, Ana Rita Dias Araújo, Aline Mairal, Akim Bello, Lucile Fleuriot, Océane Buvry, Pauline Perné, Delphine Debayle, Camille Fournes-Fraresso, Jacob B. Hansen, Pierre-Damien Denechaud, Dominique Langin, Bruno Antonny

## Abstract

Adipocytes primarily store fatty acids (FAs) as triacylglycerols (TGs) within lipid droplets, releasing them through lipolysis to meet systemic energy demands. While the metabolism of long-chain fatty acids (LCFAs) in adipocytes has been extensively characterized, it remains ill-defined how adipocytes utilize fatty acids depending on chain length and structure. Our work demonstrates that short- and medium-chain fatty acids (SMCFAs) are esterified into TGs within lipid droplets, rather than being incorporated into other FA-containing lipid species. During lipolytic activation, TGs enriched in SMCFAs are hydrolysed more rapidly than those containing LCFAs. This accelerated mobilization is facilitated by the preferential localization of SMCFA-containing TGs at the lipid droplet surface, which enhances accessibility to adipose triglyceride lipase. Unlike LCFAs, which are efficiently released for utilization by peripheral tissues, SMCFAs are predominantly oxidized within adipocytes. These findings reveal a unique metabolic routing of SMCFAs, indicating their preferential intracellular oxidation to support adipocyte energy requirements.

## Introduction

Triacylglycerol (TG) is the most abundant stored component of the white adipose tissue (WAT) and serve as the key substrate for energy mobilization^1,2^. Each TG molecule consists of a glycerol esterified by three fatty acids (FAs), which can differ in chain length (carbon number) and degree of saturation (number of double bonds). These properties shape the biophysical characteristics of TGs and other complex lipids, thereby modulating their metabolic behaviour and functions in the cell^3,4^.

FA composition of the WAT-stored TG pool is determined by three major sources: (i) FAs from diet, (ii) FAs derived from hepatic *de novo* lipogenesis, and (iii) FAs derived from WAT *de novo* lipogenesis. However, the contribution of the latter source is quantitatively minimal in humans^5^. Importantly, adipocytes can remodel FA chemistry by expressing elongase and desaturase enzymes such as ELOVL6 and SCD1^6,7^.

The esterified FA pool in WAT has been reported to be primarily composed of FAs with 16 and 18 carbon chain lengths^5^. However, FAs present in low quantities may exert significant regulatory effects on adipocyte metabolism. In particular, short- and medium-chain (<10 carbon number) FAs (SMCFAs) have distinct physical features which could influence lipid turnover, mitochondrial activity or signalling in the adipocytes^8,9^. Compared to long chain FAs (LCFAs), SMCFAs have higher water solubility which allows free transport within the cytosol and in the circulation. Additionally, unlike LCFAs – which require facilitated transport mechanisms to enter the mitochondria – SMCFAs can freely diffuse across the mitochondrial membranes, making them efficient substrates for mitochondrial β-oxidation^10,11^. While medium-chain FAs are often obtained from dietary sources, short-chain FAs are primarily produced by microbial fermentation in the gut^12,13^. Due to their rapid intestinal absorption and efficient metabolism in the liver, TGs containing medium-chain FAs serve as fast energy sources and are therefore used in patients with inborn errors of fatty acid oxidation^14,15^.

SMCFAs have been shown to trigger a wide range of biological effects in various cell types, including adipocytes^16–18^. It has been found that SMCFAs can reduce adiposity and control body weight and insulin sensitivity^19,20^. Additionally, evidence suggests that these FAs can be synthesized endogenously by human tissues as well^21^. Notably, peroxisomal shortening of LCFAs has been reported to lead to the production of SMCFAs in mammalian cells, which indicate that these FAs are endogenous members of the human lipidome and might have intrinsic functions^21–23^.

TG molecules containing SMCFAs have been recently reported in WAT^24^. However, it is not known whether they are stored within adipocytes. Furthermore, the localization and metabolism of TGs containing SMCFAs have not been investigated. To address this, we combined mass spectrometry analysis of 2D and 3D-differentiated adipocytes along with mouse models, reconstitution systems and molecular dynamics simulation. These approaches reveal that SMCFA-containing TGs show specific distribution within the lipid droplets and are used by the adipocyte itself, in contrast to LCFA-containing TGs, which are mobilized for export to other tissues. This retention of SMCFA-TGs ensures their domestic use to sustain adipocyte energy homeostasis.

## Results

### 1. SMCFAs are synthesized via *de novo* lipogenesis during adipocyte differentiation

We aimed to quantify esterified SMCFAs in human WAT. For this, we applied a three-phase lipid extraction procedure that enables the separate analysis of neutral and polar lipid fractions by LC-MS/MS^25^. Using this approach, lipids were extracted from human subcutaneous WAT samples obtained from the abdominal fat depot and proceeded for LC-MS/MS-based lipidomics. No SMCFA-containing polar lipids were identified. However, we identified 258 TG molecules of which 36 contained SMCFAs (**Fig. 1a**). Notably, in all detected SMCFA-TG species, SMCFAs were esterified at a single position, while the remaining two positions were occupied by LCFAs. Analysis of the FA composition of WAT TGs revealed that oleic acid (C18:1) and linoleic acid (C18:2) were the most abundant species. In contrast, SMCFAs accounted for only 0.2 mol% of the total TG-stored FAs. Among the identified SMCFAs, caprylic acid (C8:0) and caproic acid (C6:0) were the most abundant species (**Fig. 1b**). Interestingly, SMCFA-containing TGs are enriched in C12-16 fatty acids compared to the total TG FA pool (**Extended Data Fig. 1a**). This suggests that the incorporation of a SMCFA into DGs favours the presence of shorter FAs.

**Figure 1.**
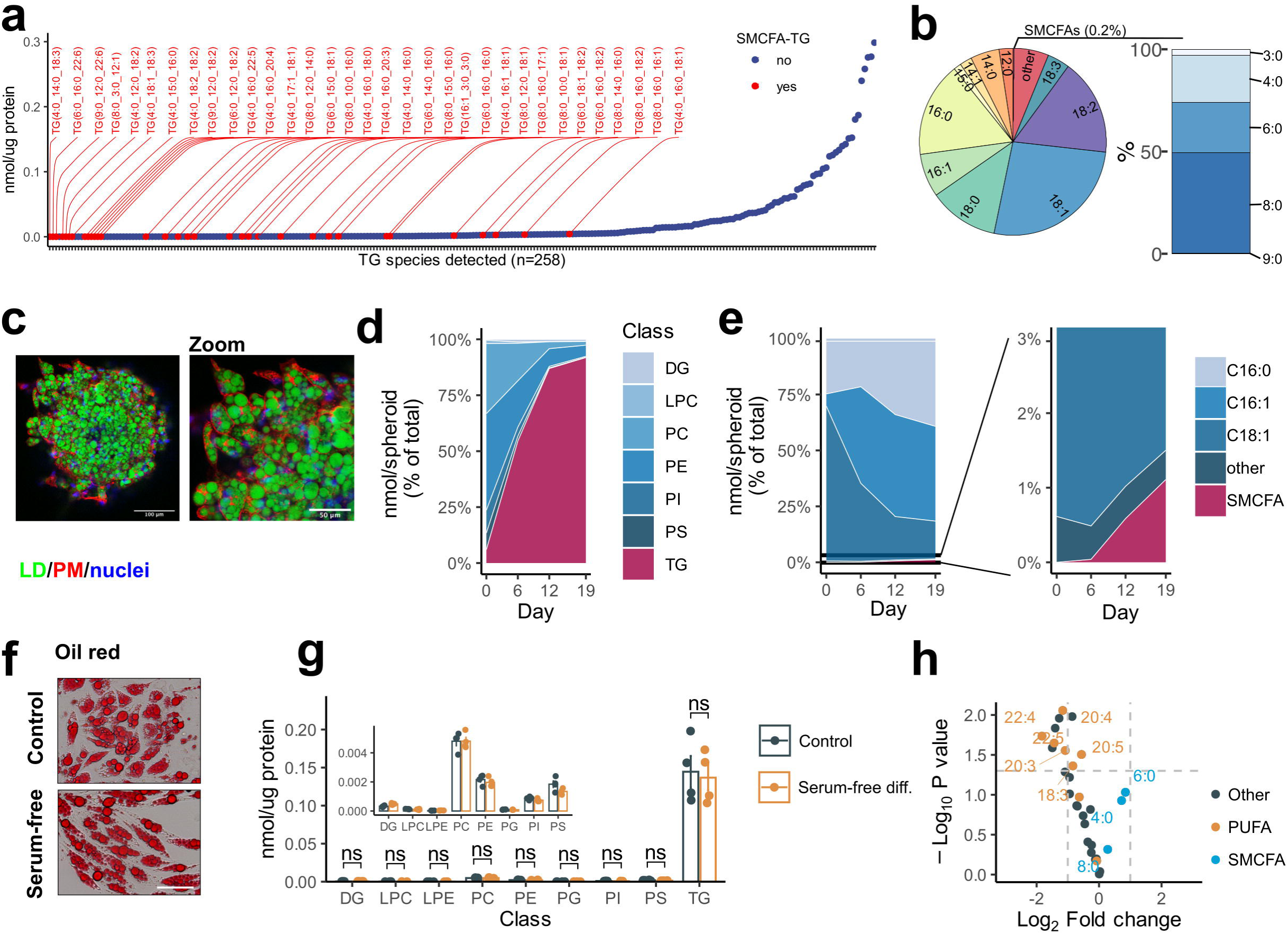
Short- and medium chain fatty acids are stored in WAT. **a-b,** Lipids were extracted from human abdominal WAT samples (*n*=5) then neutral fractions containing TG were analysed by LC-MS/MS. Protein-normalised quantities of each detected TG species (**a**). Short- and medium-chain fatty acid-containing TGs (SMCFA-TGs) are coloured red. Mean of five samples are shown. FA composition of the detected TG molecules (**b**). **c-e,** TERT-hWA pre-adipocytes were differentiated in 3D to generate spheroids. At day 19 of differentiation, spheroids were fixed, then lipid droplets (green), plasma membrane (red) and nuclei (blue) were labelled. Spheroids were visualized using confocal microscopy. Scale bars: 100 and 50 µm (**c**). TERT-hWA adipocyte spheroids were pooled at days 0, 6, 12 and 19 of differentiation, then lipid contents were analysed using LC-MS/MS. The amount of TG increased rapidly through the differentiation process and dominated the adipocyte lipidome by day 19 of differentiation (**d**). Changes of the TG-stored FA levels during TERT-hWA spheroid during differentiation. In fully differentiated spheroids, C16:0 and C16:1 were the most abundant FAs stored in TG. The proportion of SMCFAs in TGs increased progressively with differentiation. Mean values were from three biological replicates are shown. Each derived from a pool of eight spheroids. **f-h,** TERT-hWA cells were differentiated under normal or serum-free conditions, then fixed and stained with Oil Red O (**f**) or processed for lipidomics (**g, h**). Scale bar: 100 µm. **g**, no differences were detected between the intensities of the analysed lipid classes. Bars represent means ± SEM of four biological replicates. ns=non-significant, unpaired two-sample t-tests for each lipid class, followed by Bonferroni correction for multiple testing. **i**, The abundance of long, polyunsaturated FAs (PUFAs) stored in the TGs have significantly reduced upon serum-free differentiation. Log_2_ fold changes were calculated for each lipid class (*n*=4), p-values were determined using two-sample t-tests comparing the control (C) and serum-free (SF) conditions. Bonferroni correction was applied to adjust p-values for multiple testing. Adjusted p-values are shown as -log_10_P for visualization.

As different WAT depots have distinct metabolic functions, their FA composition profiles might reflect functional differences^26,27^. We compared the TG pools of paired gluteal and subcutaneous WAT (**Extended Data Fig. 1b-d**) to investigate whether they vary in SMCFA-TG compositions. Principal component analysis (PCA) of the complete lipidomics data indicates that the two human WAT depots have major differences in their overall lipidomes (**Extended Data Fig. 1b**). Pairwise comparisons and differential analysis revealed that gluteal WAT TGs contained significantly less SMCFA-TGs than the abdominal depot, indicating that SMCFAs are unequally distributed across the different WAT depots (**Extended Data Fig. 1c-d**).

We also performed quantitative lipidomics analyses on mouse perigonadal WAT. Similarly to human WAT, SMCFAs were detected only in TGs. Among the 159 TGs, 38 contained SMCFAs (**Extended Data Fig. 1e**), accounting for 1.1 mol% of the total TG-stored FA pool, with C6:0 being the most abundant (**Extended Data Fig. 1f**).

Bulk tissue analyses includes not only adipocytes, but also other cell types present in WAT, such as fibroblasts, endothelial cells, and various immune cells^28,29^. To follow the development of adipocyte lipid composition and SMCFA accumulation over time, we differentiated human WAT-derived, immortalized stromal vascular cells (TERT-hWA)^30^ into mature adipocytes and performed a time-resolved lipidomic analysis during the 19-day differentiation process. We performed cell differentiation in 3D, as it has recently been reported that such conditions result in a phenotype with morphological, transcriptomic and lipidomic features closer to intact WAT^31^. Following 19-days of differentiation, the resulting TERT-hWA spheroids contained well-differentiated adipocytes with large lipid droplets (LDs) (**Fig. 1c**). Lipidomes of the spheroids were analysed at days 0, 6, 12, and 19 of differentiation. Gradual changes in the overall spheroid lipidomes were observed throughout the differentiation process (**Extended Data Fig. 1g**) and TG became the most abundant lipid class in the fully differentiated spheroids (**Fig. 1d, Extended Data Fig.1h**). We detected a total number of 64 TG species in the differentiated spheroids, and 13 of them contained SMCFAs (**Extended Data Fig. 1i**). Using this data, we quantitatively assessed the FA compositions of the TERT-hWA spheroid TGs upon differentiation. The quantity of C16:0 and C16:1 produced through *de novo* FA synthesis was markedly increased during adipocyte differentiation (**Extended Data Fig. 1j**). Of note, SMCFAs, undetectable before the 6^th^ day of differentiation, gradually increased to represent 1 mol% of total TG-stored FA in fully differentiated adipocytes at the 19^th^ day (**Fig. 1e**).

Although we found that SMCFAs are stored in WAT, their origin remained unclear. Therefore, we determined whether SMCFAs are taken up from the environment or synthesized by adipocytes upon adipogenesis. For this, we differentiated TERT-hWA cells in control and serum-free medium and performed lipidomics analysis to compare the two lipidomes. Oil red O staining and lipid class quantifications by LC-MS/MS show that TERT-hWA cells can produce large amounts of TG and obtain a fully differentiated phenotype in the absence of serum (**Fig. 1f, g**). As expected, the amount of long, polyunsaturated FAs (PUFA) significantly dropped upon serum-free differentiation, as these FAs cannot be endogenously synthesized. In striking contrast, no significant differences were established in SMCFA levels between control and serum free differentiation, suggesting that these FAs are produced by the adipocytes via *de novo* lipogenesis (**Fig. 1h**).

We conclude that SMCFAs are produced and stored in white adipocytes and that their increasing abundance is associated with adipogenesis.

### 2. Adipocytes metabolize SMCFAs and LCFAs through distinct pathways

Because SMCFAs have physical features distinct from regular FAs, we aimed to compare the metabolism of SMCFAs and regular FAs in adipocytes. We first investigated how SMCFAs are utilized by adipocytes. To this end, we differentiated TERT-hWA adipocyte spheroids and supplemented the differentiation media with 0 or with 150 µM of C16:0, C18:1 or C6:0 from day 6 until the end of the differentiation, and then we performed lipidomics with LC-MS/MS. Next, we determined the relative changes of each detected lipid species between the control and each FA-supplemented adipocyte culture. **Fig. 2a-c** highlight the lipid species that contain C16:0, C18:1 or C6:0 among their acyl chains. C16:0 and C18:1 was detected in multiple lipid classes and became significantly more abundant in some lipid classes upon supplementations with them (**Fig. 2a-c**). In contrast, upon C6:0 supplementation, this FA was incorporated only into TG species and was not detected in other lipid classes (**Fig. 2c**). Next, we quantified the total levels of esterified C16:0, C18:1 and C6:0 upon each supplementation condition (**Extended Data Fig. 2a**). We found that C16:0 and C18:1 are highly prone to be esterified, leading to significant increases in their overall pools in various lipids. In contrast, C6:0 supplementation contributed less to its overall esterified pool (**Fig. 2d, Extended Data Fig. 2b**). Visualization of the spheroids by confocal microscopy confirmed the different storage efficacy of the three examined FAs. Spheroids treated with C16:0 or C18:1 contained enlarged LDs, whereas C6:0 supplementation did not increase LD size significantly (**Fig. 2e**). Altogether, these results suggest that the cells can store C16:0 and C18:1 more efficiently than C6:0, suggesting distinct uptake and/or utilization mechanisms.

**Fig. 2.**
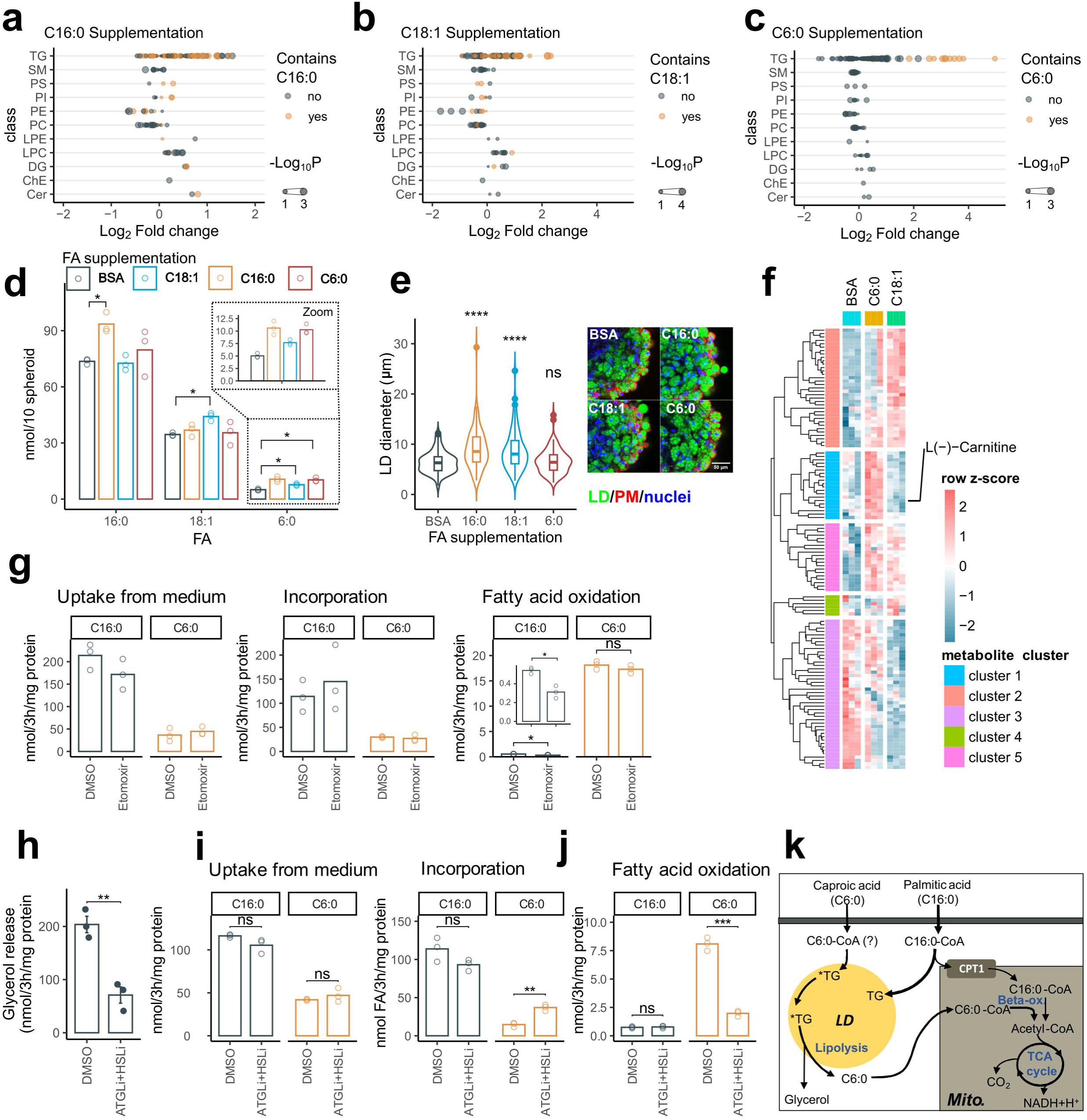
Following mobilization, adipocytes oxidize SMCFAs. **a-d,** TERT-hWA spheroids were differentiated in the presence of 30 µM BSA (Control) or 30 µM BSA complexed with 150 µM C16:0, C18:1 or C6:0. Following differentiation, lipids were extracted and analysed by LC-MS/MS. 3 biological replicates were used, each containing six pooled spheroids. **a-c**, Log_₂_ fold changes relative to the control condition were calculated for each detected lipid species and grouped by lipid class for visualization. Point size reflects −log_₁₀_ p values. Lipids containing C16:0 (**a**), C18:1 (**b**), or C6:0 (c) are highlighted; C6:0 was detected exclusively in TGs. p-values were obtained using two-sample t-tests comparing control (BSA) and fatty acid–supplemented conditions, followed by Bonferroni correction for multiple testing. **d**, Abundance of esterified C16:0, C18:1 and C6:0 in TG upon control and FA supplemented conditions. Bars represent mean values of three biological replicates. *p < 0.05, Welch’s t-test with Benjamini-Hochberg correction for multiple comparisons. **e,** Spheroids differentiated in the presence of 30 µM BSA (Control), or BSA complexed with 150 µM C16:0, C18:1 or BSA-C6:0 then fixed and stained to visualize LDs (green), Plasma membrane (PM, red) and nuclei (blue) by confocal microscopy. LD size distribution analyses revealed that C16:0 and C18:1 supplementation increased LD size. For each condition, 100 LDs were quantified from 3–4 spheroids. *p < 0.05, ****p < 0.0001, ns=non-significant, Wilcoxon rank-sum test. Scale bar: 50 µm. **f,** TERT-hWA cells were differentiated in 2D in the presence of 30 µM BSA (Control) or 30 µM BSA complexed with 150 µM C18:1 or C6:0. Following differentiations, metabolites were extracted and non-targeted metabolomics was performed using LC-MS/MS. Three biological replicates for each condition were used. Metabolites showing significant change at least once between two experimental conditions were gathered to 5 groups with hierarchical clustering. Among the metabolites upregulated upon C6:0 diet (cluster 2), we identified L-carnitine. **g,** ihASC pre-adipocytes were differentiated, then loaded with radio-labelled C16:0 or C6:0 for 3 h in the absence or presence of 50 µM CPT-1 inhibitor Etomoxir. Following labelling, FA uptake from medium, incorporation to cells and oxidation were assessed using scintillation counting of the medium, the cell lysate and the trapped CO_2_. Bars show mean values of three biological replicates. *p < 0.05, ns=non-significant, Welch’s *t*-tests. **h, i**hASC pre-adipocytes were exposed to a combination of HSL or ATGL inhibitors for 3 h (10 µM Bay and 40 µM NG-497), then lipolysis efficacy was assessed by measuring free glycerol in the cell culture supernatant. Bars show mean values ± SEM of three biological replicates. **p < 0.01, unpaired Welch’s *t*-tests. **i,** Pre-adipocytes were differentiated, then loaded with radio-labelled C16:0 or C6:0 for 3 h in the absence or presence of the HSL and ATGL inhibitor combination (10 µM Bay and 40 µM NG-497). Following labelling, FA uptake from medium, incorporation to cells and oxidation were assessed using scintillation counting of the medium, the cell lysate and the trapped CO_2_. Bars show mean values of three biological replicates. **p < 0.01, ***p < 0.005, ns=non-significant, unpaired Welch’s *t*-tests. **h**, Metabolic pathways leading to oxidation of LCFAs and SMCFAs in adipocytes. *TG: TG with C6:0.

Next, we investigated how the different FA species impact global adipocyte metabolism. We differentiated TERT-hWA adipocytes, supplemented or not the cells with C18:1 or C6:0, and then performed untargeted metabolomics. PCA of the metabolomics data revealed that the metabolome of the 3 adipocyte groups separated from each other, indicating that the incubations with the two FAs lead to different metabolomic profiles (**Extended Data Fig. 2b**). Hierarchical clustering of the altered metabolites identified compounds which are specifically enriched upon C6:0 or C18:1 diets (cluster 1 and cluster 2) (**Fig. 2f**). Importantly L-carnitine – a metabolite related to FA oxidation – was increased upon C6:0 diet, suggesting that this treatment boosts mitochondrial flux as well as energy metabolism and FA oxidation in the adipocytes.

It has been reported that SMCFAs can be rapidly oxidized in the mitochondria, however, their catabolic pathways in adipocytes have never been explored^32,33^. We tested the oxidation of C6:0 in comparison with C16:0 in another similar model of inducible human adipose stem cells (ihASC) that can readily be differentiated to adipocytes. Three hours after incubating the adipocytes with ¹UC-labeled fatty acids at the C1 position, we assessed FA uptake, cellular incorporation, and oxidation (**Extended Data Fig. 2c**). Although the uptake and the cellular incorporation levels of C16:0 were 5-times higher than that of the C6:0 (**Fig. 2g**, left and middle panel), C6:0 was much more prone to oxidation as its loading generated a 20-fold higher level of radioactive CO_2_ than C16:0 loading (**Fig. 2g**, right panel). We performed the experiments in the presence of the CPT1 inhibitor Etomoxir as well, that blocks the entry of acyl-CoAs in the mitochondria. As expected, Etomoxir treatment significantly blocked C16:0 oxidation, whereas it had no effect on C6:0 oxidation, suggesting that this FA can enter the mitochondria directly, without the CPT1 system.

Next, we aimed to test whether C6:0 is oxidized directly upon uptake or it is first incorporated into TG and stored in the LD before oxidation. As the activity of the lipolytic lipases – Adipose Triglyceride Lipase (ATGL) and Hormone Sensitive Lipase (HSL) – is a pre-requisite for the release of FAs from the LD^2^, we first tested the effects of pharmacological inhibitors targeting these lipolytic enzymes. For this, we treated adipocytes for 3 h with the combination of specific ATGL and HSL inhibitors. As reflected by the level of glycerol – the end product of lipolysis – released into the culture media^34^, this treatment reduced basal L ipolysis significantly (**Fig. 2h**). Next, we loaded adipocytes with radiolabelled C16:0 or C6:0 in the presence or absence of lipolysis inhibitors and assessed 3 h later FA uptake, incorporation and oxidation. Pharmacological lipolysis inhibition had no effect on these parameters in the case of C16:0 (**Fig. 2i**, middle and left panels). Strikingly, C6:0 oxidation almost completely abolished upon lipase inhibition (**Fig. 2i**, right panel). Furthermore, the level of C6:0 incorporation – its amount in the TG molecules – increased concomitantly upon lipolysis inhibition. These experiments indicate that, following uptake, C6:0 is incorporated into TG to be subsequently oxidized in a mechanism dependent on lipolysis. In contrast, a fraction of C16:0 can be imported directly into the mitochondria for oxidation *via* CPT1 after uptake, without requiring a prior esterification step (**Fig. 2j**).

### 3. TGs with SMCFA are highly prone to lipolysis

To better understand SMCFA-TG dynamics upon lipolysis, we analysed the TG lipidome of adipocyte spheroids under basal conditions and after stimulation with the lipolysis inducer Forskolin (FSK) (**Extended Data Fig. 3a**). As expected, three hour FSK treatment significantly increased media glycerol level, indicating an efficient stimulation of lipolysis in the spheroids (**Fig. 3a**). Following FSK stimulation, lipids were extracted from the adipocyte spheroids and the individual TG species were quantified in the neutral fractions by LC-MS/MS.

**Fig. 3.**
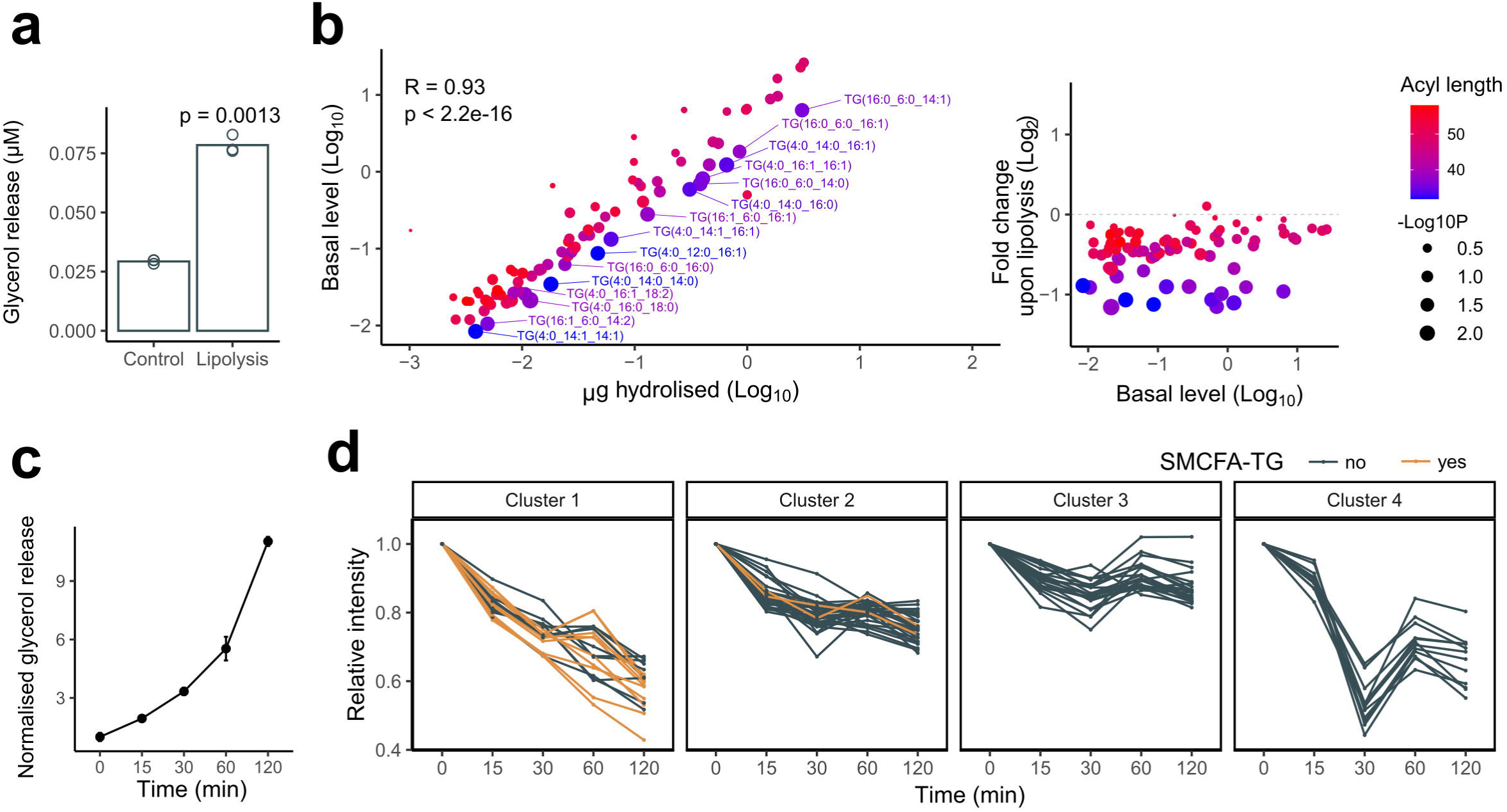
FA length drives TG lipolysis in adipocytes. **a-b,** TERT-hWA adipocytes per group was collected and incubated in the presence or absence of 1 µM Forskolin (FSK) for 3 h. Following incubation, medium was tested for glycerol release (**a**), and lipids were extracted from the spheroids for LC-MS/MS (**b**). **b**, *Left panel:* he difference between control and FSK-treated samples for each TG species (µg hydrolysed) was calculated for differential analysis. Basal levels of TG species were plotted as a function of their hydrolysed amounts upon FSK treatment. Point size reflects log-transformed p-values comparing control and FSK-treated samples, and colour denotes total acyl-chain carbon number. *Right panel*: Log_2_Fold change upon lipolysis for each TG species detected was plotted in the function of their initial amount detected in control samples. Log-transformed p-values are indicated by the size of the points, while colour represent total carbon number of the acyl chains of each TG molecule. Each condition involved three biological replicates, containing five spheroids/replicate. P-values were calculated by Welch’s t-test. **c-d,** TERT-hWA spheroids were collected in test tubes and treated with 1 µM FSK for 0, 15 30, 60 and 120 min. Following incubation, medium was tested for glycerol release (**c**) and spheroids were extracted for lipidomic analysis with LC-MS/MS. Data represent mean ± SEM from four biological replicates. Each replicated contained 10 pooled spheroid. (**b**) k-means clustering of the kinetic data indicates that SMCFA-TG species are rapidly hydrolysed upon FSK stimulation and show similar dynamics upon lipolysis. Values represent means of four biological replicates.

The extent of hydrolysis for each TG species was determined by calculating the difference between their detected amounts in the control and in the FSK stimulated spheroids (**Extended Data Fig. 3a-c**, red bars). We conducted differential analysis on the quantitative TG species data. First, we plotted the TG species abundance under basal condition against the amount that got hydrolysed upon FSK treatment **(Fig. 3b**, left panel). Overall, we observed a positive linear correlation between these two parameters indicating that TG species were globally hydrolysed during lipolysis in proportion to their abundance. However, when the data points were coloured based on the total carbon length of the fatty acids in the TG molecules, the plot reveals that the blue dots (TGs containing SMCFAs) are consistently shifted to the right compared to the red dots (TGs without SMCFAs) (**Fig. 3b**, left panel). Regression analysis revealed a statistically significant difference between the intercepts of the lines fitted to SMCFA-containing TGs and regular TGs, while their slopes did not differ significantly (**Extended Data Fig. 3d**). This right shift suggested that SMCFA-TG species are more prone to FSK-induced lipolysis compared to the other TG species with similar quantities present in the adipocyte. As a minor decrease in TG quantity could lead to high fold change values in TG species with low quantities, we tested whether the low abundance of SMCFA-TGs could create a bias in the analysis. We determined the relative susceptibility to lipolysis of all TG species (y value) against their abundance (x value) and their acyl chain length (colour code) (**Fig. 3b**, right panel). Compared to other TG species with similar abundance (similar x values), SMCFA-TGs always exhibited more susceptibility to lipolysis.

Next, we compared the lipolysis kinetics of each TG species following FSK stimulation in adipocyte spheroids (**Extended Data Fig. 3e**). Glycerol concentration in the medium showed an increase over time, indicating efficient and continuous lipolysis (**Fig. 3c**). Following neutral lipid extraction and analysis by LC-MS/MS, we plotted the relative intensity values of each TG species over time (**Extended Data Fig. 3f**). K-means clustering of the analysed TG species showed that regardless of the nature of the other two FAs present in the TG, SMCFA-TGs mostly gathered in cluster 1, which corresponds to the species that display the fastest time course of lipolysis upon FSK addition (**Fig. 3d**). Cluster 2 TGs showed a slower but continuous decay over time. Thus, the presence of SMCFA favours the breakdown of TG molecules during lipolysis. Clusters 3 and 4 contained TGs with fluctuating intensity profiles, most probably due to re-esterification and/or biosynthesis processes^35^. The mean TG acyl chain length showed a progressive increase across the four clusters, consistently rising from cluster 1 through cluster 4 (**Extended Data Fig. 3f)**.

Altogether, these findings indicate that TGs containing SMCFAs are highly susceptible to lipolysis in adipocytes.

### 4. TGs with SMCFAs accumulate at the LD surface

A possible explanation for the observed preference of lipases towards SMCFA-TG is that the distribution of the two applied TG species differs within the LDs according to FA chain length. If TGs with SMCFAs, being more polar, are positioned at the surface of LDs, they may be more accessible to lipases, thereby resulting in a faster hydrolysis rate.

To test this hypothesis, we first investigated lipolysis dynamics *in vitro* in minimalistic systems and modelled the behaviour of LDs containing SMCFA-TGs. First, we prepared artificial lipid droplets (aLDs) composed of either TGs with three oleic acids – TG(18:1/18:1/18:1) – or three hexanoic acids – TG(6:0/6:0/6:0), respectively. We produced aLDs with phosphatidylcholine (PC) monolayers to mimic the phospholipid fraction of the LDs^36^. Following aLD preparations, we used pancreatic lipase to induce lipolysis and aLDs incubated with heat-inactivated lipase served as control. Following incubation with the lipase preparations, lipids were extracted and the quantities of the TG species were monitored in each sample using LC-MS/MS. Upon lipase heat inactivation, the TG species were readily detected (**Fig. 4a**). The active lipase induced a comparable reduction of the TG signals for both substrates, indicating that the lipase hydrolysed aLDs containing each TG species with similar efficiency (**Fig. 4a**). To investigate lipase preference for TGs when both substrates are present in the LDs, we prepared aLDs containing a 5:95 (mol:mol) ratio of TG(6:0/6:0/6:0) to TG(18:1/18:1/18:1), mimicking their physiological proportions. As shown in **Fig. 4b**, the lipase almost completely hydrolysed the TG(6:0/6:0/6:0) pool (95% hydrolysis), while the hydrolysis of TG(18:1/18:1/18:1) was modest (20% hydrolysis), indicating that when the two TGs are present in the same LDs, TG with SCFAs are hydrolysed with higher efficacy than LCFAs TGs. This selective hydrolysis efficacy is further supported by fold change analysis of the intensity values between active and inactive lipase conditions: TG(6:0_6:0_6:0) exhibited a markedly higher drop in fold change than TG(18:1_18:1_18:1), confirming the preferential hydrolysis of the SMCFA-TG by the lipase.

**Fig. 4.**
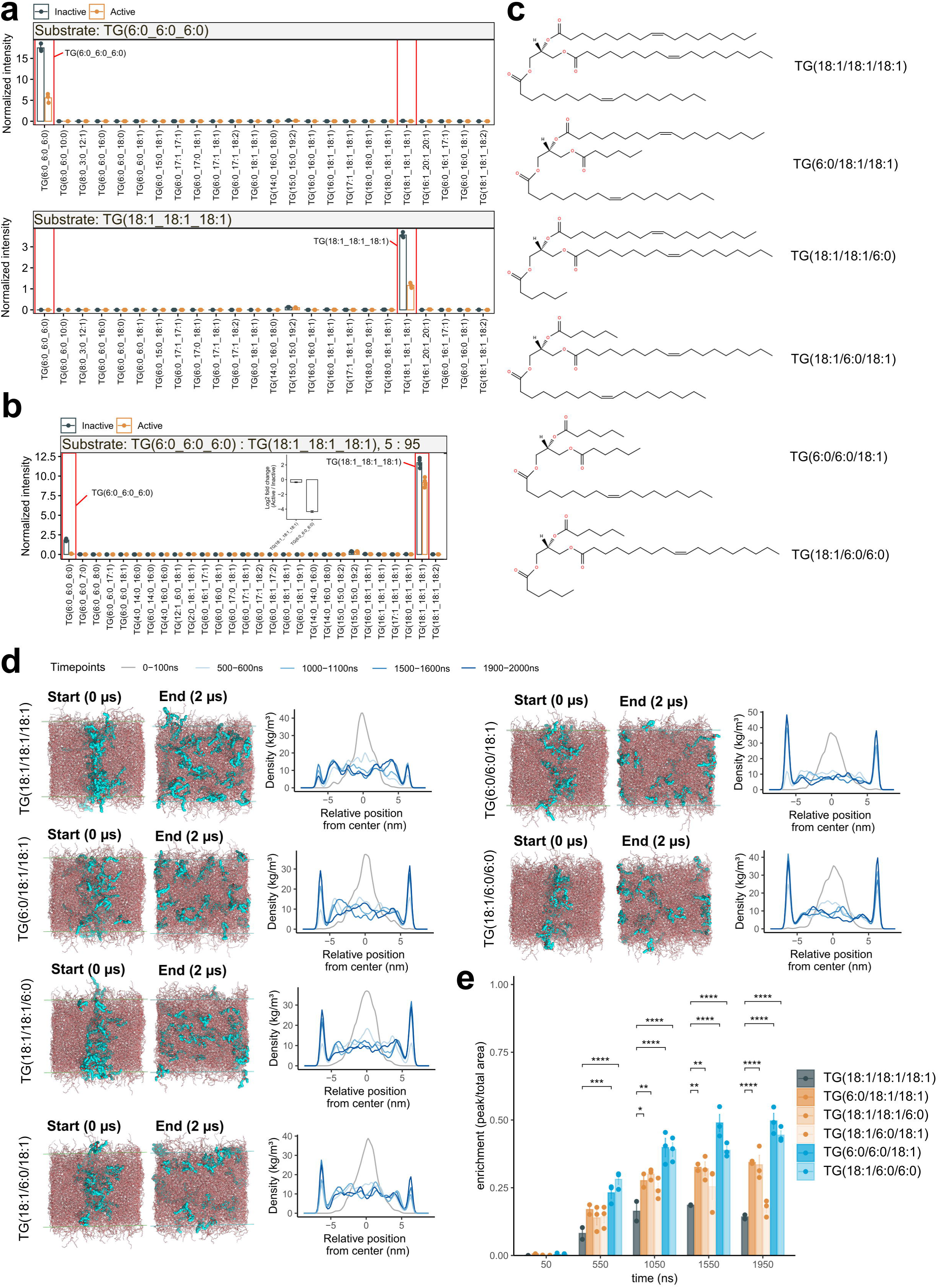
Short-chain FAs directs TG molecules to the LD surface. **a-b,** Artificial lipid droplets composed of TG(6:0_6:0_6:0), TG(18:1_18:1_18:1) or a 5:95 molar ratio mixture of the two were prepared and incubated with active or heat-inactivated pancreatic lipase. Following incubations, neutral lipids were extracted and analysed by LC-MS/MS. Lipase activity on each substrate alone was comparable (**a**). When both substrates were present, TG(6:0_6:0_6:0) was preferentially hydrolysed as indicated by the larger Log_2_ fold change value corresponding to this substrate (inlet) (**b**) Bars indicate mean ± SEM values calculated from three technical replicates. The experiment was repeated three times with similar results. **c,** Chemical structures of triolein (TG(18:1/18:1/18:1)) and the custom TG species containing C6:0 fatty acids used in molecular dynamics simulations. **d–e**, In the simulations, 5% of the total TG molecules were replaced with one of the test species and positioned at the central plane of the simulation box. Over a 2Uµs time frame, the spatial distribution of test TGs was monitored. TGs containing C6:0 at the sn-1 and sn-3 positions accumulated at the oil–water interface, whereas triolein showed no such positioning. This interfacial enrichment was enhanced when both sn-1 and sn-3 positions contained C6:0. (**e**) Bar plots show the relative enrichment of test molecules at the oil–water interface at different time points. Bars represent meanU±USEM from 2-3 independent simulations. Statistical comparisons based on estimated marginal means (EMMs) from a linear model using triolein as the reference. Bonferroni correction was applied for multiple comparisons. Significance levels are indicated as follows: *pU<U0.05, **pU<U0.01, ***pU<U0.001 and ****pU<U0.0001.

As an alternative approach, we next performed molecular dynamics simulations with TG(18:1/18:1/18:1) and custom constructed SMCFA-TG species TG(6:0/18:1/18:1) to better understand the positioning of SMCFA-TGs to the LD surface. We constructed and tested the three possible stereoisomers by placing the hexanoic acid to either the sn-1, 2 or to the sn-3 positions of the glycerol backbone (**Fig. 4c**). Recent studies indicate that the LD surface is not completely covered by phospholipids and surface-oriented TG molecules can create an interface which promotes protein recruitment^37,38^. Therefore, molecular dynamics simulations were conducted within a lateral box containing water-TG interfaces at the left and right edges. The vertical dimensions of the box were set to be infinitely extended to simulate bulk-like conditions. 5% of the total TG pool was assigned as test molecule and positioned to the middle vertical plane of the simulation box (Start time). The distribution of these test molecules was monitored throughout a 2 µs simulation period, then their lateral density was calculated at the indicated time points (**Fig. 4d**). Subsequently, relative accumulations at the interfaces were quantified (**Fig. 4e**).

TG(18:1/18:1/18:1) did not exhibit specific accumulation at the interfaces during the simulation (**Fig. 4d, e**). In contrast, TG(6:0/18:1/18:1) molecules, where the 6:0 fatty acid esterified at either the sn-1 or sn-3 position, began to accumulate at the water-TG interface shortly after the simulation started. Interestingly, this surface accumulation effect was less pronounced when C6:0 was esterified at the sn-2 position of the TG molecule (**Fig. 4d, e**, orange columns). Next, we conducted similar experiments with TGs containing two 6:0 and one 18:1 FAs (**Fig. 4d, e**). These lipids started to accumulate on the surface rapidly and reached a maximal enrichment at the interfaces shortly after launching the simulations (**Fig. 4e**, blue columns). This enrichment was higher than that observed with TG with only one short chain 6:0 FA.

From the *in vitro* lipolysis experiments and molecular dynamic simulations, we conclude that SMCFA-TGs are more effectively targeted by lipases compared to regular TGs because of their preferential segregation at the surface of the LDs.

### 5. ATGL is the enzyme responsible for release of SMCFAs stored in TGs

Surface exposure of SMCFA-TGs raises the question of the involvement of the two adipocyte neutral lipases, ATGL and HSL. ATGL hydrolyses the first ester bond of the TG molecules to release one FA molecule and produce diacylglycerol (DG)^39,40^. HSL is the sole DG hydrolase in adipocytes, however, it also possesses TG hydrolase activity^41^.

To test that SMCFA-TGs are hydrolysed by these canonical lipolytic enzymes, we used a tamoxifen-inducible CreER^T^^2^/LoxP mouse model to generate primary adipocytes deficient in both ATGL and HSL. Pre-adipocytes were isolated from WAT stromal vascular fractions of mice carrying floxed alleles of *Pnpla2* (ATGL) and *Lipe* (HSL) and expressing or not the inducible CreER^T^^2^ recombinase. Adipocyte differentiation was initiated *in vitro*, then two days later, gene deletion was induced by 4-hydroxy-tamoxifen treatment (**Extended Data Fig. 4a**). Western blot and RT-qPCR data show that the expression levels of both ATGL and HSL were successfully reduced by the treatment in the primary adipocyte cultures expressing the recombinase (**Fig. 5a-b**). Importantly, loss of the two lipolytic enzymes did not impair adipocyte differentiation, as comparable lipid accumulation levels were detected in control and double knock-out cells (**Fig. 5c**). Lipolysis was induced in the primary adipocytes with the β_3_-adrenergic receptor agonist CL316343. Measurement of glycerol levels in the media indicated that both basal and stimulated lipolysis were reduced in the neutral lipase-deficient adipocytes (**Fig. 5d**). In parallel, cells were harvested to assess the hydrolysis of SMCFA-TGs by LC-MS/MS. In control adipocytes, SMCFA-TGs were highly prone to lipolysis as reported above in TERT-hWA spheroids (**Fig. 3**, **Fig. 5e**, left panel). In neutral lipase-deficient adipocytes, TG hydrolysis – including SMCFA-TGs – was less prominent than in the control cells (**Fig. 5e**, right panel). Both inhibition of the lipase catalytic activity in human adipocytes (**Fig. 2h, i**) and decreased expression of the enzymes in mouse adipocytes (**Fig. 5d, e**) indicate that the canonical pathway involving ATGL and HSL contribute to the lipolysis of SMCFA-TGs.

**Fig. 5.**
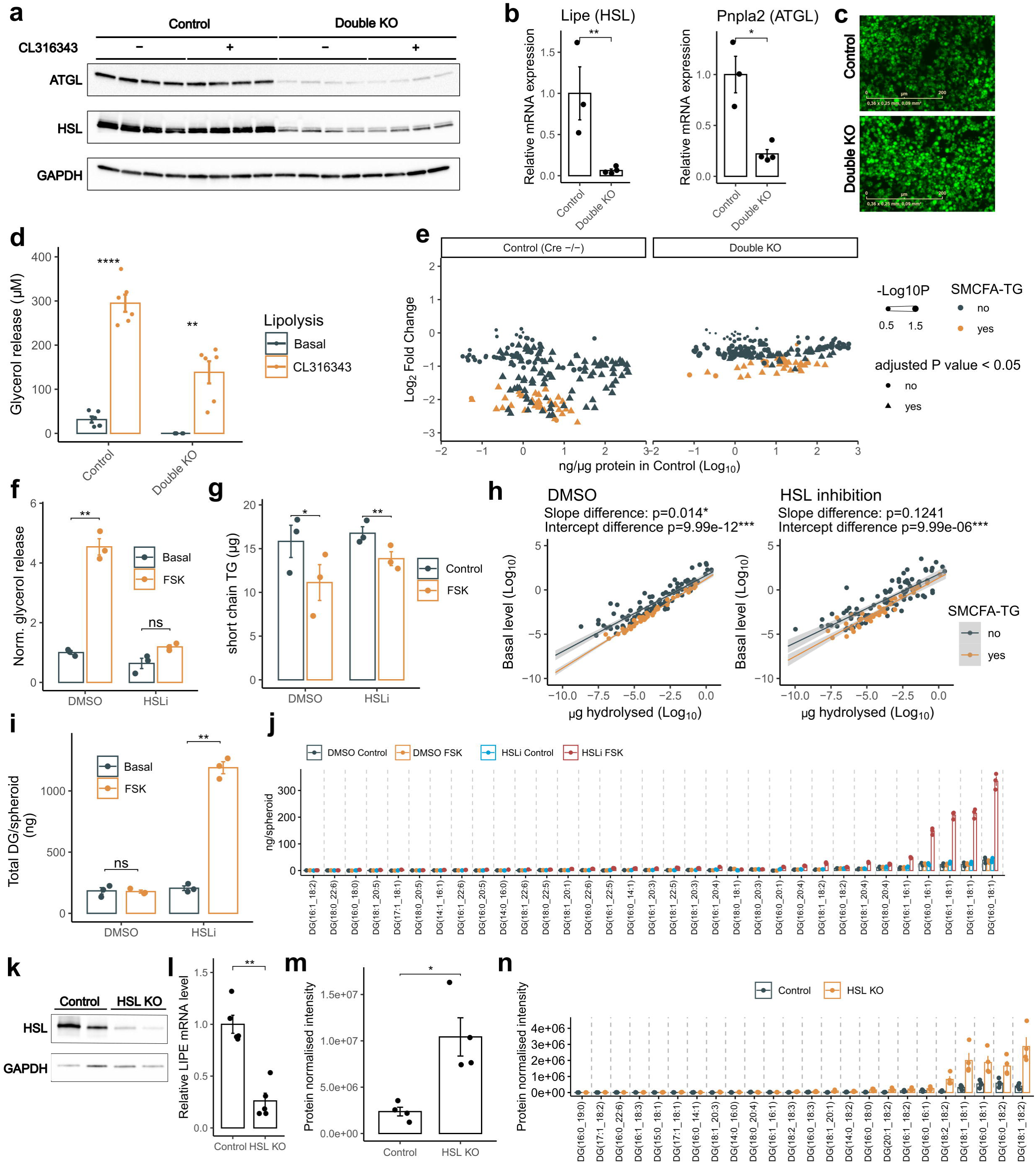
ATGL targets TGs with short- and medium-chain FAs. **a-e,** Stromal vascular cells were isolated from 2-5 day old AdipoCre+ and AdipoCre- mice carrying floxed *Lipe* (HSL) and *Pnpla2* (ATGL) alleles. Following *in vitro* adipogenic differentiation, 4-hydroxy-tamoxifen was applied to activate the recombinase and generate Double KO cells. Cells were stimulated with 1 µM CL316343 for 2 h, then the cells were collected for western blot (**a**), and RT-qPCR (**b**) to validate efficient gene depletion. Bars indicate mean ± SEM of 3-4 animal/group, *p<0.05, **p<0.05, Unpaired t-test. **c**, representative fluorescent micrographs of Control and Double KO cells stained with BODIPY indicate no marked differences in lipid accumulation between the two conditions. Scale bar: 200 µm. **d**, glycerol was measured in the medium following lipolysis induction. Bars represent mean ± SEM values calculated from six biological replicates, **p<0.01, ****p<0.01, Welch’s t-test with Bonferroni correction. **e,** Neutral lipids were extracted from Control and Double KO primary adipocyte cultures following lipolysis inductions then LC-MS/MS was performed. Log_2_ Fold change values between lipolysis and basal conditions were calculated for each detected TG molecule, then plotted against their initial quantity in basal conditions. In Double KO cells, overall efficacy of TG lipolysis including SMCFA-TGs was reduced. Four biological replicates/experimental group was used. P values were calculated by Welch’s t-test with Bonferroni correction. **f-j,** TERT-hWA spheroids were treated for 3 h with DMSO or 1 µM Forskolin (FSK) in the presence or absence of HSL inhibitor (10 µM Bay). Following treatment, glycerol levels were measured in the supernatant (**f**), and lipids were extracted from the spheroids for LC-MS/MS analysis. Total levels of SMCFA-TGs dropped significantly upon FSK stimulation upon both control and HSL inhibitor treatments (**g**). Differential lipid species analysis between basal and stimulated conditions was performed for each treatment. FSK stimulation preferentially enhanced the hydrolysis of SMCFA-TGs in both DMSO- and HSL inhibitor-treated spheroids. Regression lines were fitted to points corresponding to SMCFA-TG and regular TGs, then statistical differences in their slopes and their intercepts were established. No significant difference between the intercepts of the regression lines was detected. Slopes were compared by linear model ANOVA and intercepts were tested by coefficient t-tests. Bars represent the meanU±USEM of three biological replicates, each consisting of a pool of 8–10 spheroids (**h**). FSK increased total DGs in HSL-inhibited spheroids (**i**). Quantities of each detected DG species. No DG with SMCFA were detected (**j**). Statistical comparisons were performed by two-way ANOVA test between basal and stimulated conditions within each inhibitor group followed by a Bonferroni correction for multiple testing. Significance is indicated as: *pU<U0.05, **pU<U0.01. **k-n,** AdipoCre+ and AdipoCre-(Control) mice carrying floxed *Lipe* (HSL KO) allele were fed with 4-hydroxy-tamoxifen to induce adipocyte-specific depletion of HSL. Following gene depletion, WAT was collected and specific HSL depletion was confirmed with western blot (**k**) and RT-qPCR (**l**). For RT-qPCR measurements, five animals were used for each experimental group. Following gene depletions, WAT samples were collected and lipids were extracted to perform LC-MS/MS. Compared to the control, significant increase in total DG level was detected in the adipose tissue of HSL KO mice (**m**). Protein-normalised intensity values of all DG species detected (**n**). Bars represent meanU±USEM. On **m-n,** Four mice/group was analysed. *p<0.05, **p<0.01, ns=non-significant, Unpaired t-test.

Next, we decided to identify which of these two lipases mediates the hydrolysis of SMCFA-TGs. Notably, no DG species with SMCFAs were detected in our previous lipidomics experiments (**Fig. 1a-e**), suggesting that these FAs are preferentially removed during the initial step of lipolysis. We therefore induced lipolysis in TERT-hWA spheroids using FSK in the absence or presence of a pharmacological HSL inhibitor and subsequently analysed changes in TG and DG species abundance. As expected, FSK-stimulated glycerol release was blocked by HSL inhibition, consistent with the role of this enzyme in converting DG to monoacylglycerol (**Fig. 5f**). Importantly, the loss in the total amount of SMCFA-TGs was preserved upon HSL inhibition despite blockade of the hydrolysis of two ester bonds (**Fig. 5g**). We determined the changes in abundance of all detected TG species upon lipolysis in DMSO and HSL inhibitor-treated adipocytes. In DMSO-treated control spheroids, comparison of the fitted linear models to regular TGs and SMCFA-TGs revealed a significant difference in their intercepts. This vertical shift between the regression lines aligns with our previous results (**Fig. 3** and **Fig. 5e**) and confirms that SMCFA-TGs are more susceptible to lipolysis than other TG species (**Fig. 5h**, left column). Importantly, in the presence of the HSL inhibitor, SMCFA-TGs remained the most affected species, as evidenced by a significant difference in the intercepts of the two fitted regression lines (**Fig. 5h**, right panel). Aligning with the role of HSL as a DG hydrolase, significant accumulation in DG was observed when lipolysis was stimulated and HSL inhibited (**Fig. 5i**)^42^. No SMCFAs were detected in the DG accumulated as a result of HSL inhibition (**Fig. 5j, Extended Data Fig. 4b, c**).

To confirm the contribution of ATGL activity on SMCFA-TG hydrolysis *in vivo* we investigated WAT in mice lacking HSL in adipocytes by LC-MS/MS. Following 4-hydroxy-tamoxifen treatments to activate the CreER^T2^ recombinase in mice carrying floxed *Lipe* (HSL) allele, HSL was successfully depleted in the WAT (**Fig. 5k, l**). WAT samples derived from HSL knock-out mice showed a massive accumulation of DG compared to control WAT samples (**Fig. 5m**). Notably, despite this DG accumulation, no SMCFA-containing DG species were detected (**Fig. 5n, Extended Data Fig. 4d**).

Together, these results indicate that ATGL actively hydrolyses SMCFA-containing TG molecules.

## Discussion

Due to recent developments of mass spectrometry-based lipid analysis, a more detailed view on the chemical composition of the WAT lipidome is now emerging^45,46^. Next to bulk lipid species such as long-chain fatty acid-containing phospholipids and TGs, recent research has uncovered a growing number of rare and non-abundant lipid species with distinct metabolic functions^1,24,47,48^. Among them, SMCFAs represent a unique class as they exhibit distinct biophysical properties that may determine their metabolic routes as compared to other FA species. Lipidomic profiling revealed the presence of triglycerides containing SMCFAs in human and mouse WAT^24,47^. Here, we show that FAs with short- and medium carbon length acyl chains (C4-10) are endogenously synthesized to be stored in adipocyte LDs and demonstrate that these FAs are readily mobilized to be oxidized within adipocytes.

Our data demonstrate that SMCFA-TGs exhibit enhanced susceptibility to lipolysis compared to their long-chain FA containing counterparts. This elevated lipolytic susceptibility was consistently observed across multiple experimental systems, including human adipocyte spheroids, primary mouse adipocytes and *in vitro* LD reconstruction models. Using molecular dynamic simulations, we observed that SMCFA-containing TGs preferentially accumulate at the LD surface, especially when the 6:0 fatty acid occupies the sn-1 or sn-3 positions. By demonstrating how FA chain length and positional isomerism influence TG localization within the LD, our findings highlight that LD architecture is a critical regulator of substrate selection during lipolysis^49^. This spatial organization represents a novel mechanism by which adipocytes prioritize the mobilization of specific lipid species to efficiently meet metabolic demands.

The active zone of a lipid droplet corresponds to the thin layer of lipids and proteins that form its surface, where all metabolic reactions occur. The thickness of this active zone (L) should be in the range of approximately 2 nm, roughly the dimension of a TG or a phospholipid molecule. Compared to the bulk volume of a droplet (V_t_ = 4/3πR^3^), the volume of the active zone is given by V_a_= 4πR^2^L, yielding a ratio of active-to-bulk of V_a_/ V_t_ = 3L/R. For a 100 nm LD, this fraction is in the range of 6%. However, for a large LD as that found in white adipocyte (R = 10 - 100 µm), V_a_/ V_t_ = 0.06 - 0.006%. This means that even minor TG species, such as SMCFA-TGs, can have a significant impact on the surface reactivity of the lipid droplet, provided they preferentially partition into the active zone.

We concluded that the preferential hydrolysis of SMCFA-TGs is dependent on ATGL activity. By removing one FA from the TG molecule, ATGL catalyses the initial and rate-limiting step of lipolysis, while HSL primarily converts this DG to monoacylglycerol^40–42^. Stimulated lipolysis experiments on 3D adipocyte cultures treated with HSL inhibitor confirmed this observation: when ATGL remained active, the selective SMCFA-TG hydrolysis was still detectable despite the overall reduced efficiency of the stimulated lipolysis. This strongly indicates that ATGL is the principal enzyme targeting SMCFA-TGs in adipocytes. Notably, upon our lipidomics analyses we did not detect DGs with SMCFAs, supporting the notion that ATGL preferentially cleaves the short-chain fatty acid directly from TGs. However, we cannot exclude the possibility that this absence is due to the detection limits of our mass spectrometry approach.

Our results show that adipocytes can synthesize SMCFA-TGs *de novo* during differentiation, without relying on external FA sources. Intriguingly, chronic supplementations of LCFAs lead to a concurrent increase in esterified SMCFAs levels in adipocytes, suggesting a potential link between LCFA oxidation and SMCFA biosynthesis. Although the precise steps of the SMCFA biosynthetic pathways remain to be elucidated, one possibility is that LCFAs can undergo incomplete oxidation, which could lead to the release of shortened FAs from mitochondria. However, we should not exclude the possibility that SMCFAs originate from non-mitochondrial FA oxidation or are directly synthesized from acetyl-CoA. Such pathways may include fatty acid synthase-dependent premature chain termination or the activity of short- and medium-chain thioesterases, both of which require further exploration^50–52^.

The fate of SMCFAs after their mobilization is also distinct from that of LCFAs. We revealed that SMCFA, such as C6:0, are preferentially oxidized by adipocytes compared to LCFAs. SMCFAs undergo significantly higher rates of mitochondrial oxidation, bypassing the canonical CPT1-dependent mitochondrial transport system. It should be noted that incorporation of SMCFAs into phospholipids would compromise membrane stability and lipid packing. Thus, oxidation provides an efficient route for adipocytes to rapidly dispose these FAs, preventing their accumulation in membrane lipids to maintain lipid homeostasis. In addition, oxidation of SMCFAs depends on basal lipolytic activity, as inhibition of lipases reduced their catabolism, indicating that stored SMCFAs are mobilized from lipid droplets before oxidation. This metabolic routing suggests a tightly coupled system in which SMCFA-TGs are selectively hydrolysed and rapidly oxidized, providing a prompt and efficient energy source for the adipocyte.

Taken together, our findings reveal a distinct class of FAs with specialized metabolic roles in adipocytes. The fatty acid chain length governs their intracellular fate. TGs containing SMCFAs preferentially localize at the lipid droplet surface, rendering them highly susceptible to lipolysis. Upon hydrolysis, LCFAs are typically released, whereas SMCFAs are selectively retained and rapidly oxidized within the cell to supply immediate energy. Our findings position FA chain length as a new determinant of lipid remodelling in adipocytes, complementing recent insights into TG cycling and adipose storage dynamics^35,53^. Further research is needed to explore whether dysregulation of SMCFA metabolism occurs in metabolic disorders such as obesity and type 2 diabetes.

## Methods

### 1. Adipocyte differentiation, spheroids and FA supplementations

TERT-hWA pre-adipocytes were maintained in Advanced DMEM-F12 medium (Gibco, 12634-028) completed with 10% FBS (Biowest), 2 mM L-Glutamine (Gibco, 25030-024) 100 U/mL Penicillin and 100 µg/mL Streptomycin (Gibco, 15140-148) and 2.5 ng/mL β-FGF (Sigma-Aldrich, F3685). For differentiations in 2D, medium was replaced to 2% FBS culture medium on confluent TERT-hWA cell, then 2 days later cells were primed with adipogenic cocktail (1 nM 3,3’,5-Triiodo-L-thyroine (Sigma-Aldrich, T6397), 5 µg/mL insulin (Sigma-Aldrich, I6634), 1 µM Rosiglitasone (Sigma-Aldrich, R2408), 1 µM hydrocortisone (Sigma-Aldrich, H0369), 1 µM Dexamethasone (Sigma-Aldrich D4902), 0.5 mM 3-Isobutyl-1-methylxantine (Sigma-Aldrich, I5879)) for 6 days. Following this, medium was replaced to 2% FBS containing culture media and refreshed at every 3 days. Cells were considered fully-differentiated at least 13 days following adipogenic priming. For differentiations in 3D spheroids, pre-adipocytes were seeded into ultra-low attachment plates (Greiner bio-one) in 5000 cells/well density, then centrifuged at 150 x g for 2 minutes to promote spheroid formation. 4 days later, medium was replaced to adipogenic priming medium, then changed 6 days later to 2% FBS containing medium for at least 13 days. Differentiation was confirmed by microscopic evaluation of HCS LipidTOX^TM^ Green Neutral Lipid Stain (ThermoFisher Scientific), Alexa Fluor 594-conjugated Wheat Germ Agglutinin (ThermoFisher Scientific) and Hoechst 33342 (ThermoFisher Scientific)-labelled spheroids using a confocal microscope (Leica, TCS-SP5) equipped with a HCX IRAPO L 25×/0.95 water objective. LD diameters were measured in ImageJ.

For different fatty acid supplementations (C6:0-hexanoic acid (Sigma-Aldrich, 153745), C16:0-palmitic acid (Sigma-Aldrich, P0500) and C18:1Δ9-oleic acid (Sigma-Aldrich, O1383), FAs were conjugated with fatty acid-free BSA (Sigma-Aldrich, 12657) in 1:5 BSA-FA molar ratio^54^, then conjugates were applied on the adipocytes following adipogenic priming in 150 µM FA concentrations.

ihASC pre-adipocytes from AstraZeneca were cultured in Endothelial Cell Growth Medium MV2 Kit supplemented with 5% FCS and growth factors (PromoCell, C22121), and completed with 1% Penicillin-Streptomycin. After confluence, cells were switched to the differentiation medium (Zen Bio, BM-1) completed with 3% FBS, 1% Penicillin-Streptomycin and adipogenic cocktail: 100 nM human insulin (Sigma-Aldrich, I9278), 1 µM Dexamethasone, 0.5 mM 3-Isobutyl-1-methylxantine, 1 µM Pioglitazone (Sigma-Aldrich, E6910) for the 6 first days. Following that, 3-Isobutyl-1-methylxantine and Pioglitazone were omitted. The medium was refreshed at every 2-3 days until complete cell differentiation at day 12.

### 2. Animal studies and primary cells

The mouse models used in this study were generated previously by insertion of LoxP sites in the *Lipe* gene encoding HSL^42^, or in the *Pnpla2* gene encoding ATGL^55^. To obtain adipocyte specific HSL knock-out, *Lipe^fl/fl^* mice were crossed with transgenic mice (Tg(AdipoQ-Cre/ERT2)1Soff/J) expressing the tamoxifen-inducible Cre/ERT2 recombinase, expressed under the control of the adiponectin promoter, specific for adipose tissue^56^. 7-week old mice were gavaged with tamoxifen (1 mg/day for 5 days, Sigma-Aldrich, T5648) to induce adipocyte-specific deletion of HSL (HSL KO). Control littermate were *Lipe*^fl/fl^ AdipoQ-Cre/ERT2^-^ mice. Mice were fed standard chow (Ssniff V1534) and housed according to Inserm guidelines and European Directive 2010/63/UE in the local animal care facility (agreements A 31 555 04 and C 31 555 07). Protocols were approved by the French Ministry of Research after review by local ethical committee (Comité d’éthique en experimentation animale de l’UMS006/CREFRE, CEEA122, Toulouse, France). Blood was collected in EDTA-coated tubes (Sarstedt) through tail vein sampling. Plasma was obtained after centrifugation by collecting the supernatant. Serum glycerol and non-esterified fatty acid levels were measured by colorimetric assays (F6428 Sigma and 91696 Wako).

Primary adipocytes were obtained according to the protocol described earlier^57^ from *Lipe^fl/fl^, Pnpla2^fl/fl^, AdipoQ-Cre/ERT2^-^* (Control*) and Lipe^fl/fl^ Pnpla2l^fl/fl^ AdipoQ-Cre/ERT2^+^* (Double KO) mice. Briefly, following isolation from adipose tissue, stromal vascular fraction was plated. After proliferation, pre-adipocytes were differentiated 2 days using 170 nM Rosiglitazone (Cayman Chemical, CAY-7140-100), 500 µM 3-Isobutyl-1-methylxantine, 1 µM dexamethasone and 170 nM insulin in DMEM 1 mM glucose medium (Sigma-Aldrich, D5546) supplemented with 10% FBS. The deletion of HLS and ATGL was induced by the addition of 4-hydroxy-tamoxifen between day 2 and 4 post differentiation (Sigma-Aldrich, H6278). After 7 days, the adipocytes were differentiated and used for the experiments. To estimate primary adipocyte differentiation, fixed cultures were labelled with BODIPY and visualised by an epifluorescent microscope.

### 3. Oil Red O staining

Following 2D differentiations, adipocytes were washed twice with PBS then fixed with 4% paraformaldehyde. Following two washing steps with PBS, cells were treated with 60% isopropanol, then stained with Oil Red O (Sigma-Aldrich, O0625, 1.8 mg/mL in 60% isopropanol) for 5 minutes. Cells were washed two times with tap water and imaged in a Cytation5 Cell imaging multimode-reader.

### 4. Stimulated lipolysis

For lipolysis assays, 5-8 spheroids/sample were pooled into plastic test tubes in 2% FA-free BSA containing, Advanced DMEM-F12 medium. Upon experiments performed in 2D-differentiated adipocytes, culture medium was replaced to 2% FA-free BSA containing, phenol-red free DMEM medium. Lipolysis was stimulated with 1 µM Forskolin (Sigma-Aldrich, F3917) treatment for indicated incubation times at 37°C. Following Forskolin treatments, media was harvested and glycerol production was measured by a fluorimetric assay (Sigma-Aldrich, MAK-117). For primary cells, following a 24 h starvation without serum and insulin, lipolysis was stimulated for 2 h with the β3-adrenergic receptor agonist CL316343 at 100 nM (Sigma-Aldrich, C5976).

### 5. Human tissues

WAT samples were obtained from five healthy female donors (mean age ± SD, 36.6 ± 8.6 years; BMI ± SD, 26.3 ± 1.4 kg/m²). Tissue samples were collected from both abdominal and gluteal depots during elective surgical procedures. All participants provided written consent prior to inclusion in the study. Sample collection and use for research were approved by the appropriate institutional ethics committee.

### 6. RNA isolation, RT-qPCR and Western blot

Adipose tissues were homogenized in Qiazol buffer (Qiagen) using precellys tissue homogenizer. Adipocytes were lysed in RLT buffer (Qiagen) supplemented with ß-mercaptoethanol (Sigma) accordingly to supplier. After purification on RNeasy column (Qiagen), RNA was reverse-transcribed using Multiscribe Reverse Transcriptase (Applied Biosystemts) and quantified via qPCR using QS5 instrument (Applied Biosystemts). Relative gene expressions were determined using delta Ct method normalized by mouse RS9 and 36B4 housekeeping mRNA quantifications. The primer sequences used are: Atgl (5’-TCCTTAGGAGGAATGGCCTAC-3’ and 5’-TCCTCTTCCTGGGGGACAAC-3’), Hsl (5’-ACTTCTGGAAAGCCTTCTGGAACA-3’ and 5’-GATGCCATGTTGGCCAGAGAC-3’), Rs9 (5’-CGGCCCGGGAGCTGTTGACG-3’ and 5’-CTGCTTGCGGACCCTAATGTGACG-3’) and 36B4 (5’-AGATTCGGGATATGCTGTTGG-3’ and 5’-TCGGGTCCTAGACCAGTGTTC-3’).

For immunoblot, total proteins were extracted from adipose tissues or adipocytes using protein lysis buffer (50mM Tris-HCL pH 7-8, 5 mM EDTA, 150 mM NaCl, 1% Triton-X-100 with protease and phosphatase inhibitors). After quantification with BCA assay, protein samples were resolved on SDS-PAGE and transferred to a nitrocellulose membrane. Immunoblot was revealed using ECL and imaged with a ChemiDoc software (Biorad). The following antibodies obtained from Cell Signaling Technology were used with the appropriated secondary antibodies: ATGL (n°2138), LHS (n°18381), GAPDH (n°2118).

### 7. Lipid extractions for LC-MS/MS

The following reagents were used to lipid extractions: Acetonitrile (ACN), chloroform (CHCl_3_), methanol (MeOH) and hexane (Hex) were HPLC grade; ethyl acetate (VWR, 23880.290, EtAc), butylhydroxytoluene (Sigma-Aldrich, 34750, BHT). Lipid standards (reference, abbreviation) were acquired from Sigma-Aldrich and Avanti Research: Splash lipidomix (330707, Splash), glycerol trihexanoate (T0888, TG6:0/6:0/6:0), 1,3-dipentadecanoyl-2-oleyol(d7)-glycerol (791648, dTG), 1-pentadecanoyl-2-oleyol(d7)-sn-glycerol (791647, dDG), 1-oleoyl(d7)-rac-glycerol (791646, dMG).

To extract lipids from cells and spheroids, a 3-phase liquid extraction was performed to separate neutral and polar lipids^25,58^. Adipocytes were collected in a minimum phosphate-buffered saline (PBS) volume (aqueous). Cells or spheroids were transferred into glass tubes, containing already the internal standard mix (Splash) and TG 6:0/6:0/6:0. Then, to reduce sample loss, 0.75 mL of ACN were added to microtubes, vortexed, and transferred into the corresponding glass tubes. The remaining solvents were added to each sample: 0.75 mL Hex, 0.25 mL EtAc. The aqueous phase (sample) was completed with ultrapure water when needed, resulting in Hex:EtAc:ACN:Aqueous (3:1:3:2, V:V:V:V). 2D cultured samples were vortexed and centrifuged at 2500 g, for 4 min, at 18°C. Spheroid samples were first vortexed for 30 min at 4°C with glass beads, and only then centrifuged. The upper phase was collected into a new tube with a Hamilton glass syringe, which was washed three times in solvent between samples. Hex was added (half the volume of the first extraction) to the 2 remaining phases for re-extraction. Samples were again vortexed and centrifuged, and upper and middle phases were collected separately. To reduce phospholipid loss, a re-extraction of middle phase was done with ACN:EtAc (3:1, V:V) (half the volume of first extraction). Extraction (solvents and water) and experiment blanks (PBS) were included. All extraction solvents had 50 μg/mL BHT. Extracted samples were kept dried at -20°C under Ar.

For tissue and 2D culture samples, a fixed volume of aqueous phase was collected and dried under vacuum centrifugation. RIPA buffer was added to the samples, and they were vortexed and then centrifuged at 14,000 rpm, 5 min, at 4°C. Soluble protein was quantified using a Pierce BCA kit (ThermoScientific, A55864).

### 8. Metabolite extraction for LC-MS/MS

Adipocytes were re-suspended in methanol/water, 80:20, v/v, vortex homogenized for 1 min, then samples were incubated at −20 °C for 30 minutes. This vortexing/freezing cycle was repeated three times to optimize metabolite extraction. Following the final cycle, samples were centrifuged at 3,000 rpm for 5 minutes at 4 °C. The remaining protein pellet was washed with acetone and extracted with RIPA buffer, then protein concentration was determined using a Pierce BCA Protein Assay Kit (ThermoFisher Scientific, Waltham, MA, USA). The supernatant was collected and the solvents were evaporated to dryness. Dried extracts were reconstituted in 100 µL of water/acetonitrile (90:10, v/v) containing 0.1% formic acid. After centrifugation (12,000 rpm, 15 minutes, 4 °C), 90 µL of the supernatant were transferred into LC vials and stored under an argon atmosphere until LC-MS/MS analysis.

### 9. Metabolomics by LC-MS/MS

Metabolite separation was performed by reversed-phase chromatography on a UPLC system (Ultimate 3000, ThermoFisher Scientific) equipped with a Synergi Hydro RP column (250 × 2 mm, 4 µm; Phenomenex). Chromatography was conducted at 40°C with a flow rate of 300 µL/min. The injection volume was 5 µL. Mobile phases consisted of water with 0.1% formic acid (solvent A) and acetonitrile with 0.1% formic acid (solvent B). The elution gradient (%B) was programmed as follows: 0.0 min, 0%; 7.0 min, 0%; 18.0 min, 98%. The UPLC system was coupled to a Q-Exactive mass spectrometer (ThermoFisher Scientific, San Jose, CA, USA) equipped with a heated electrospray ionization (HESI) source and operated in both positive and negative ion modes. Instrument control and data acquisition were performed using Xcalibur software (version 4.1.31.9).

Data were acquired in full MS and data-dependent MS/MS (dd-MS^2^) modes. Full MS spectra were collected at 70,000 resolution (at m/z 200) across a mass range of m/z 60–900, with an AGC target of 1 × 10^6^ and a maximum injection time of 100 ms. The five most intense precursor ions were isolated (1.5 m/z window) and fragmented by higher-energy C-trap dissociation (HCD) at a normalized collision energy of 30 eV. MS/MS spectra were acquired at 17,500 resolution (at m/z 200), with an AGC target of 1 × 10^5^ and a maximum injection time of 50 ms.

Raw data were processed using Compound Discoverer 3.3 (ThermoFisher Scientific). The untargeted metabolomics workflow was applied, including peak extraction, alignment (retention time tolerance 0.2 min; mass tolerance ±5 ppm), and peak detection (minimum intensity 50,000). Elemental composition constraints were set to: C 90, H 190, O 18, N 5, S 5, P 3, Cl 4. The final dataset (retention time–m/z pairs, annotations, name/formula, and peak areas) was exported as an Excel file for subsequent analysis.

### 10. Lipidomics by LC-MS/MS

Lipidomic profiling was performed using reverse-phase ultra-performance liquid chromatography (RP-UPLC) coupled with High Resolution mass spectrometry. Lipid extracts were separated on an ultimate 3000 (ThermoFisher Scientific, USA) equipped with an accucore C18 150x2.1, 2.5 µm column (ThermoFisher Scientific). The column was maintained at 35°C and the mobile phase was delivered at a flow rate of 400 µl/min. The injection volume was set to 3 µL. The mobile phases consisted of (A) Acetonitrile/Water 50/50 (V/V) with 10 mM ammonium formate and 0.1% formic acid and (B) Isopropanol/Acetonitrile/Water 88/10/2 (V/V) containing 2 mM ammonium formate and 0.02 % formic acid. The elution was carried out using the following stepwise gradient of solvent B: 0.0 min, 35%; 4.0 min, 60%; 8.0 min, 70%; 16.0 min, 85%; 25.0 min, 97%.

The UPLC system was coupled with a Q-exactive Orbitrap Mass Spectrometer (ThermoFisher Scientific) equipped with a heated electrospray ionization (HESI) source. Data acquisition was conducted in positive ionization mode using Xcalibur software (v4.1.31.9). Full scan MS spectra were acquired in data-dependent MS2 (dd-MS2) mode at a resolution of 70 000 (at m/z 200) over a mass range of m/z 250-1200. The automatic gain control (AGC) target was set to 1 × 10U with a maximum injection time of 100 ms. The 15 most intense precursor ions per sac cycle were selected for fragmentation. Precursor isolation was performed with a 1.0 m/z isolation window, and fragmentation was achieved using higher-energy collisional dissociation (HCD) at normalized collision energy (NCE) of 25 and 30 eV. MS/MS spectra were acquired in the ion trap at a resolution of 35,000 (at m/z 200) with an AGC target of 1 × 10U and a maximum injection time of 80 ms.

Raw data were reprocessed using LipidSearch 4.2.21 (ThermoFisher Scientific). Lipid identification was performed in product search mode, based on accurate mass and MS/MS fragmentation patterns. The following parameters were applied: precursor ions mass tolerance of 5 ppm, product tolerance 5 ppm; ion and adducts searched [H+], [NH4+] and [Na+]; alignment retention time tolerance 0.4 min; ID quality filters A, B and C.

### 11. Fatty acid oxidation assay

Fatty acid oxidation measurements were performed as previously described with minor modifications^59^. Cells were incubated in glucose-free DMEM (Gibco, 11966-025) completed with 1 mM L-carnitine, 12.5 mM HEPES, 0.25% free-fatty acid BSA and 80 µM final non-labeled palmitate or hexanoic acid supplemented with [1-¹UC]-palmitate (0.1 µCi/µL, Revvity) or [1-¹UC]-hexanoic acid (0.1 µCi/µL, American Radiolabeled Chemicals). For inhibitor treatments, the medium was completed with 50 µM Etomoxir (Sigma, 236020), or the combination of 10 µM HSLi (BAY 59-9435) and 40 µM ATGLi (NG-497, Cayman Chemical, 2598242-66-9)^34^. Adipocytes were treated with 0.5 mL of the prepared pre-heated medium in 12-well plates, sealed with Parafilm, and incubated at 37°C for 3 hours. To measure FA oxidation, 400 µL of the medium was transferred to the bottom wells of CO_₂_-capture Teflon plates, and 200 µL of 1 N NaOH was added to the top wells. Plates were sealed, and 40 µL of 70% perchloric acid was injected into the bottom wells using a Hamilton syringe. After 1 hour of agitation at room temperature, the NaOH from the top wells was transferred to scintillation vials, and 3 mL of liquid scintillation cocktail (Revvity, 6013389) was added prior to counting. Total activity controls (10 µL of medium before and after incubation with the cells) were included in order to determine FA uptake. To measure FA incorporations to the cells, cells were rinsed twice with 0.5 mL DPBS, lysed in 200 µL of 0.1% SDS and radioactivity was measured in 10 µL lysate. Scintillation counting was performed in a Tri-Carb 2100TR Liquid Scintillation Analyzer instrument (Revvity). For normalizations, protein concentrations were measured using the BCA Protein Assay (Pierce) with standards prepared in 0.1% SDS.

### 12. Artificial lipid droplets and *in vitro* lipolysis

Lipids (reference, abbreviation) were acquired from Sigma-Aldrich and Avanti Research: the coating lipid was 1-palmitoyl-2-oleoyl-glycero-3-phosphocholine (850457, POPC) and the core lipids were glycerol trihexanoate (T0888, TG 6:0/6:0/6:0) and glycerol trioleate (T7140, TG 18:1/18:1/18:1). aLDs with a theoretical diameter of 2 μm, with a 0.8% oil in lipase buffer proportion were prepared according to the calculations (eq. 2) and general method described previously ^38^. aLDs were prepared with each separate core oil, and with 95:5 molar mixtures of TG 18:1/18:1/18:1 (major) with TG 6:0/6:0/6:0.

Lipid extraction of aLD-lipase reactions: A single extraction step of 100 µL of sample was performed using the Bligh & Dyer ratios^60^. Total volumes were 1000, 1000 and 900 µL of MeOH, CHCl_3_ and aqueous fraction (water+sample), respectively. Deuterated TG, DG, and MG were used as internal standards.

### 13. Molecular dynamics simulation

All-atom MD simulations were performed with GROMACS 2021.3^61^ (https://www.gromacs.org/) with the Charmm36 forced field^62^. The triolein (TOGL - 18:1/18:1/18:1) topology was modified as Campomanes et al., 2021. The topologies of triglycerides with short chains (TOS1 - C6:0/C18:1/C18:1, TOS2 - C18:1/C6:0/C18:1, TOS3 - C18:1/C18:1/C6:0, TOD1 - C6:0/C6:0/C18:1, TOD3 - C18:1/C6:0/C6:0) has been created from the TOGL topology. The system is composed with several layers of Triolein (864 molecules) with a layer of solvent molecules (TIP3P waters and 120mM NaCl) 2.5nm thick on each side. We applied a home-made python script to randomly select 5% of TOGL (43 lipids) at the centre of the lipids layer and changed to triolein with short chain (TOSx or TODx). We applied the same method to rename 5% of TOGL to follow these lipids during the simulations as a control. A cut-off distance of 1.2 nm was used for generating the neighbour list and this list was updated at every step. Long-range electrostatic interactions were calculated using the particle mesh Ewald summation methods (PME). Periodic boundary conditions were used. During the production run, the V-rescale thermostat and Parrinello–Rahman barostat stabilized the temperature at 310 K and pressure at 1 bar, respectively. The simulations were performed for 2 µs for each system and coordinates were saved every 100 ps. 3 replicas have been performed for each system. The MD analyses of density profiles were performed using GROMACS utilities.

### 14. Lipidomics and metabolomics data analysis, statistics

Data processing was performed in R (v4.4.2.) using RStudio (v2024.12.1.563.) and dplyr and tidyr packages. Raw lipidomics data were processed by converting intensity values into µg and nmol values for each lipid species using the internal standard intensities. For tissue samples and 2D-differentiated adipocytes, concentrations were normalized to total protein content determined from the aqueous phase, while spheroid data were normalized to the number of spheroids extracted. Molar amounts of individual esterified FAs were calculated from the molar quantities of the lipid species containing them, summed, and expressed as a percentage of the total esterified FA pool. Principal component analysis (PCA) of metabolomics and lipidomics datasets using the protein-normalised values was conducted using the factoextra R package.

Differential analysis of the protein-normalised intensity values obtained to the metabolomics data was performed by DESeq2 package in R. Those metabolites which showed significant differences at least in one comparison were clustered with hierarchical clustering and plotted with pheatmap.

For differential lipid species analysis, log_2_ fold-change and –log_10_P values were computed, with P-values adjusted using the Bonferroni method by base R. Lipolysis dynamics of regular and SMCFA-containing TGs were assessed by fitting linear regression models; then differences in slopes and intercepts between groups were evaluated by comparing models with and without interaction terms with ANOVA.

For lipolysis kinetics analysis, intensity values were normalized to the t = 0 time point, scaled using the matrixStats R package, and clustered into four groups using k-means clustering implemented in the stats package.

Plots were generated using the ggplot2, ggpubr and ggsci packages. Statistical tests applied to compare experimental groups are specified in the respective figure legends.

## Author contributions

*DK* conceptualized the work, performed the experiments with the TERT-hWA adipocytes, analysed the data and wrote the manuscript. *RG* performed the molecular dynamic simulation tests. *ARDA* prepared aLDs, performed *in vitro* lipase tests, DLS measurements, optimized and performed lipid extractions from all sample types. *AB* helped with adipocyte spheroid preparations. *LF, OB, PP* and *DD* performed metabolite extractions and performed the LC-MS/MS analyses. *AM* assisted to the AstraZeneca adipocyte differentiations and performed the fatty acid oxidation measurements. *CFF* and *PDD* maintained mice, collected adipose tissues from mice and performed the lipolysis experiments on primary cells. *PDD* supervised the experiments performed on mice and primary mouse adipocytes. *JBH* provided TERT-hWA cells. *DL* provided human adipose tissue samples, supervised the experiments performed on mice and primary cells and directed the fatty acid oxidation measurements and wrote the manuscript. *BA* conceptualized the work, supervised the project and wrote the manuscript.

## Acknowledgement

This work was supported by the European Research Council (856404—SPHERES). *DK* was supported by the János Bolyai Research Scholarship of the Hungarian Academy of Sciences (BO/00019/25/8). Financial support was also provided by the Incubation Competence Centre of the Centre of Excellence for Interdisciplinary Research, Development and Innovation of the University of Szeged. *PDD* was supported by EFSD-Boehringer Ingelheim European Research Programme on “Multi-System Challenges in Diabetes, Obesity and Cardiometabolic DIsease”. We are grateful to Mikael Ryden, Niklas Mejhert (Karolinska Institutet, Sweden) and Takeshi Harayama (CNRS-IPMC, France) for their valuable feedback and help with the manuscript preparations.

## Data availability

All data used for the evaluation of this study are provided in the corresponding source files. Raw mass spectrometry files are going to be deposited to the Metabolomics Workbench repository with the PR002673 Project number (DOI: 10.21228/M84P1D).

## Code Availability

R scripts used for data analysis are available upon request from the corresponding authors.

## Extended Data Figure Legends

**Extended Data Figure 1.**
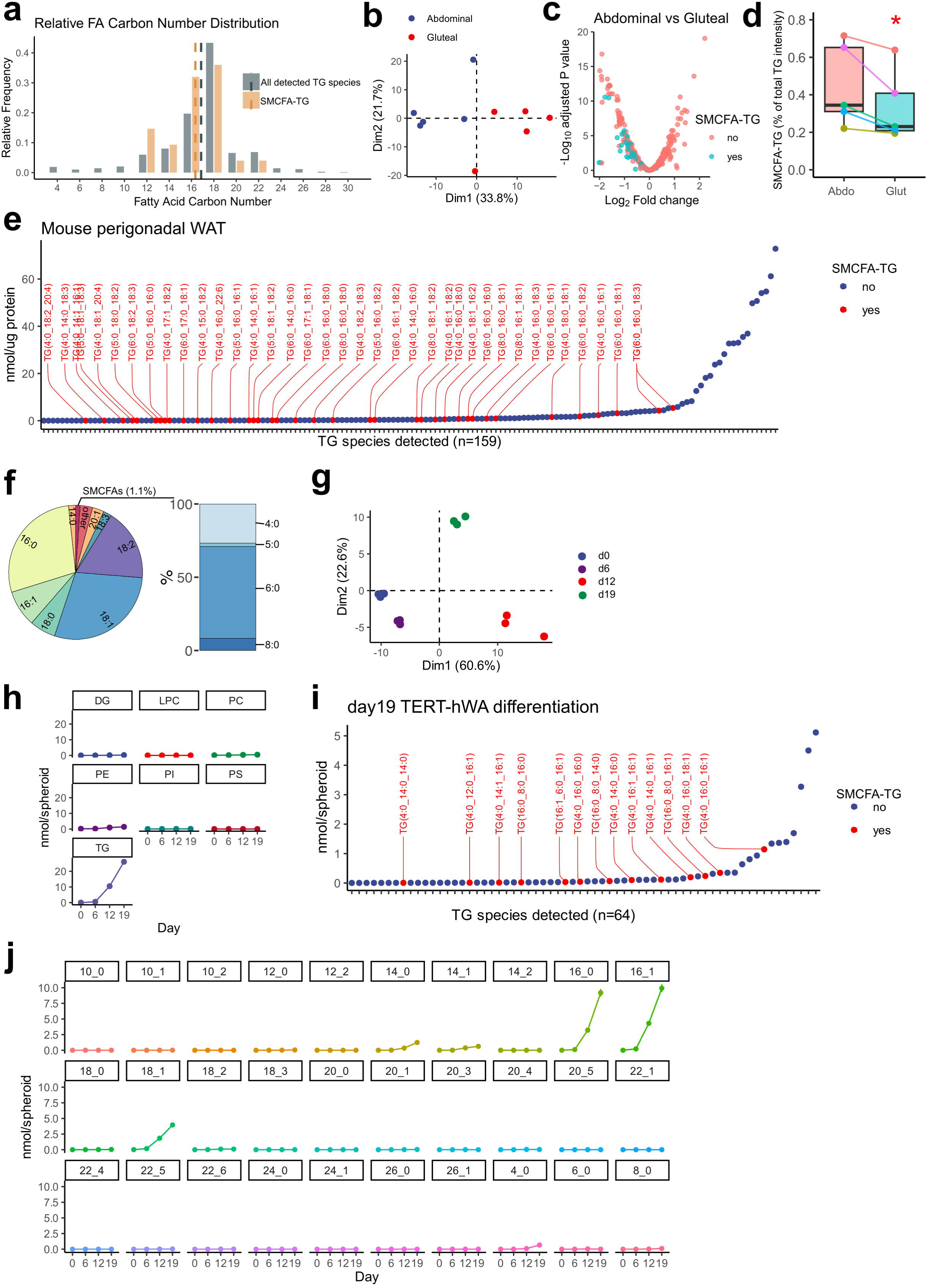
**a,** Distribution of fatty acids with even-numbered carbon chains in total TGs and in SMCFA-containing TGs. Compared to all TGs, the mean carbon length of the regular chain fatty acids (non-SMCFA) in SMCFA-TGs is shorter. **b-d**, PCA analysis of the lipidomics data obtained from paired abdominal and gluteal human WAT samples. Differential analysis of the paired abdominal and gluteal WAT samples. Compared to the abdominal depots, most SMCFA-TGs are less abundant in gluteal fat. P-values were calculated with Welch’s t-test with Bonferroni correction for multiple testing (**c**). Compared to paired abdominal WAT samples, the total amount of SMCFA-TGs was lower in gluteal WAT depots. Paired samples were obtained from five individuals. \**p*= 0.0309, Paired t-test (**d**). **e-f**, Lipids were extracted from mouse perigonadal WAT samples (*n*=4) then neutral fractions containing TG were analysed by LC-MS/MS. Protein-normalised quantities of each detected TG species (**e**). Mean of four samples are shown. SMCFA-TGs are coloured red. FA composition of the detected TG molecules (**f**). **g-j,** PCA analysis of adipocyte spheroids over time highlights progressive remodelling of the lipidome during differentiation (**g**). Abundance of each lipid class over the adipocyte spheroid differentiation, with points indicating the mean of three biological replicates (**h**). Quantity of all TG species detected in 19 day differentiated adipocyte spheroids. SMCFA-TGs are highlighted with red. Mean of three biological replicates are shown (**i**). Abundance of each esterified FA level stored in TG over the time course of the adipocyte differentiation (**j**). Points are indicating the mean of three biological replicates ± SEM.

**Extended Data Figure 2.**
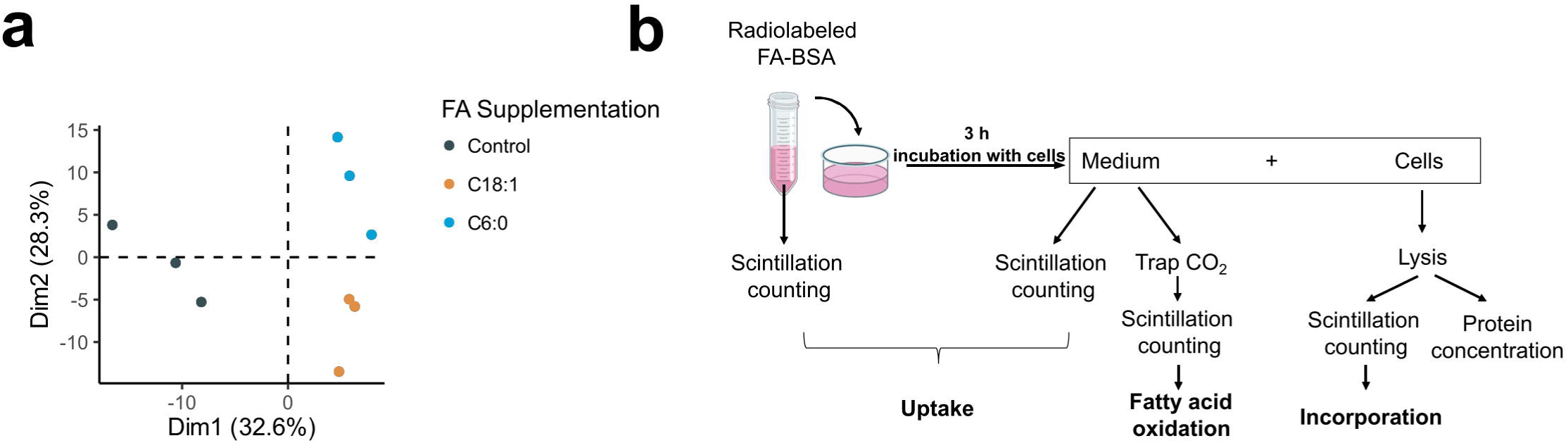
**a,** PCA analysis of the metabolome data shown on **Fig. 2f**. **b**, Schematic drawing of the fatty acid oxidation assays using radiolabelled FAs.

**Extended Data Figure 3.**
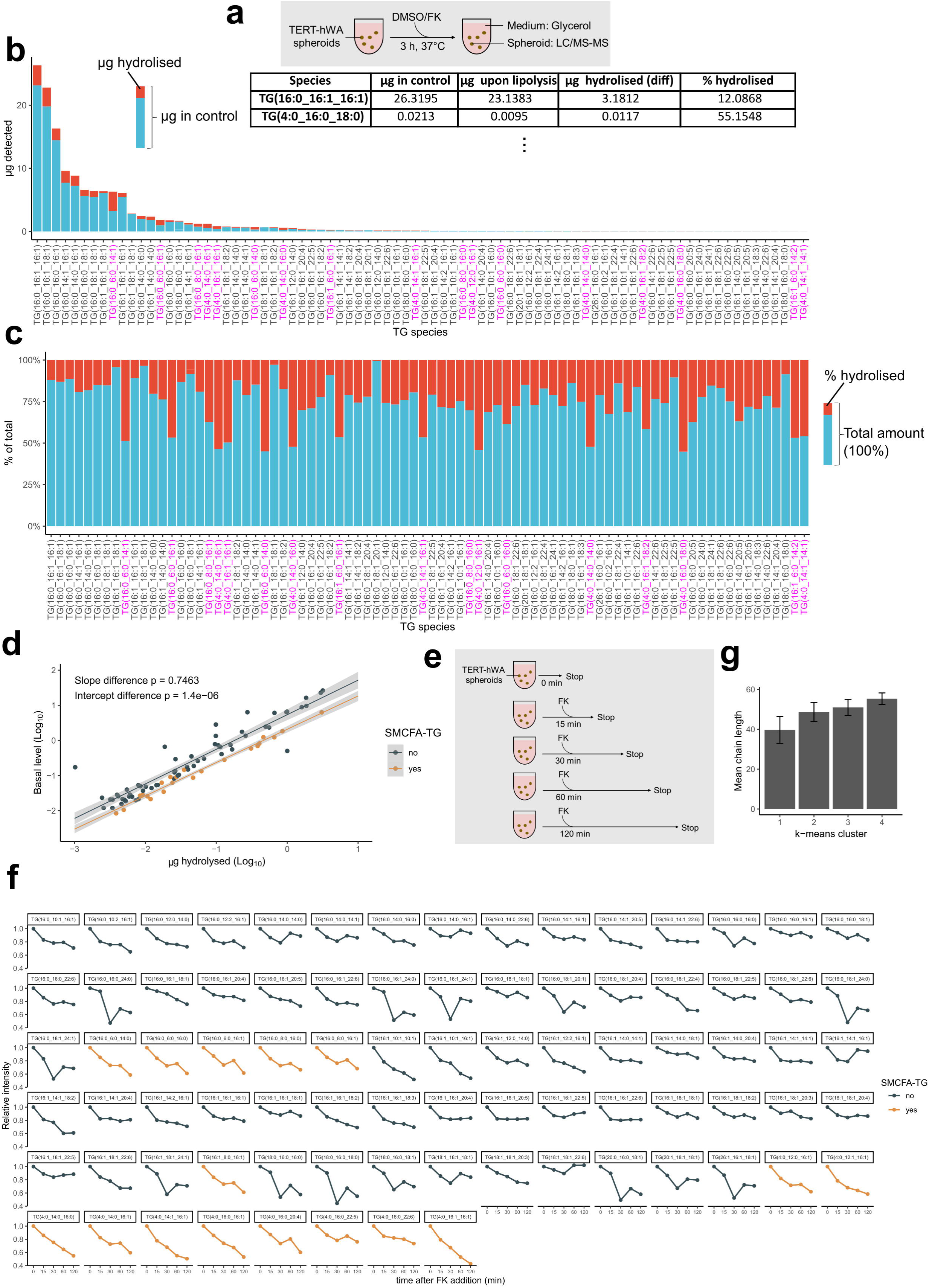
**a,** Schematic representation of the stimulated lipolysis experiments using adipocyte spheroids. Example data illustrating the calculation methods used to obtain values shown on **Fig. 3b**. **b-c**, Red bars indicate the proportion of each TG molecule that was hydrolysed following the stimulated lipolysis (**b**). Red bars depict the percentage of hydrolysis for each TG molecule under stimulated lipolysis (**c**). Three biological replicates were used to calculate TG species quantities in each condition. **d**, Basal TG levels versus hydrolysed amounts. Separate linear models were fitted for regular TGs and SMCFA-TGs. Slope differences were tested by ANOVA; intercepts by coefficient t-tests. Significant intercept difference indicates a right shift of SMCFA-TGs. **e**, Schematic representation of the stimulated lipolysis time-course in adipocyte spheroids. **f**, Changes of each detected TG species over time. SMCFA-TGs are highlighted. Means of three biological replicates are shown. **g**, Mean acyl-chain length of the TG molecules in each k-means cluster shown on **Fig. 3d**. Bars represent mean ± SEM.

**Extended Data Figure 4.**
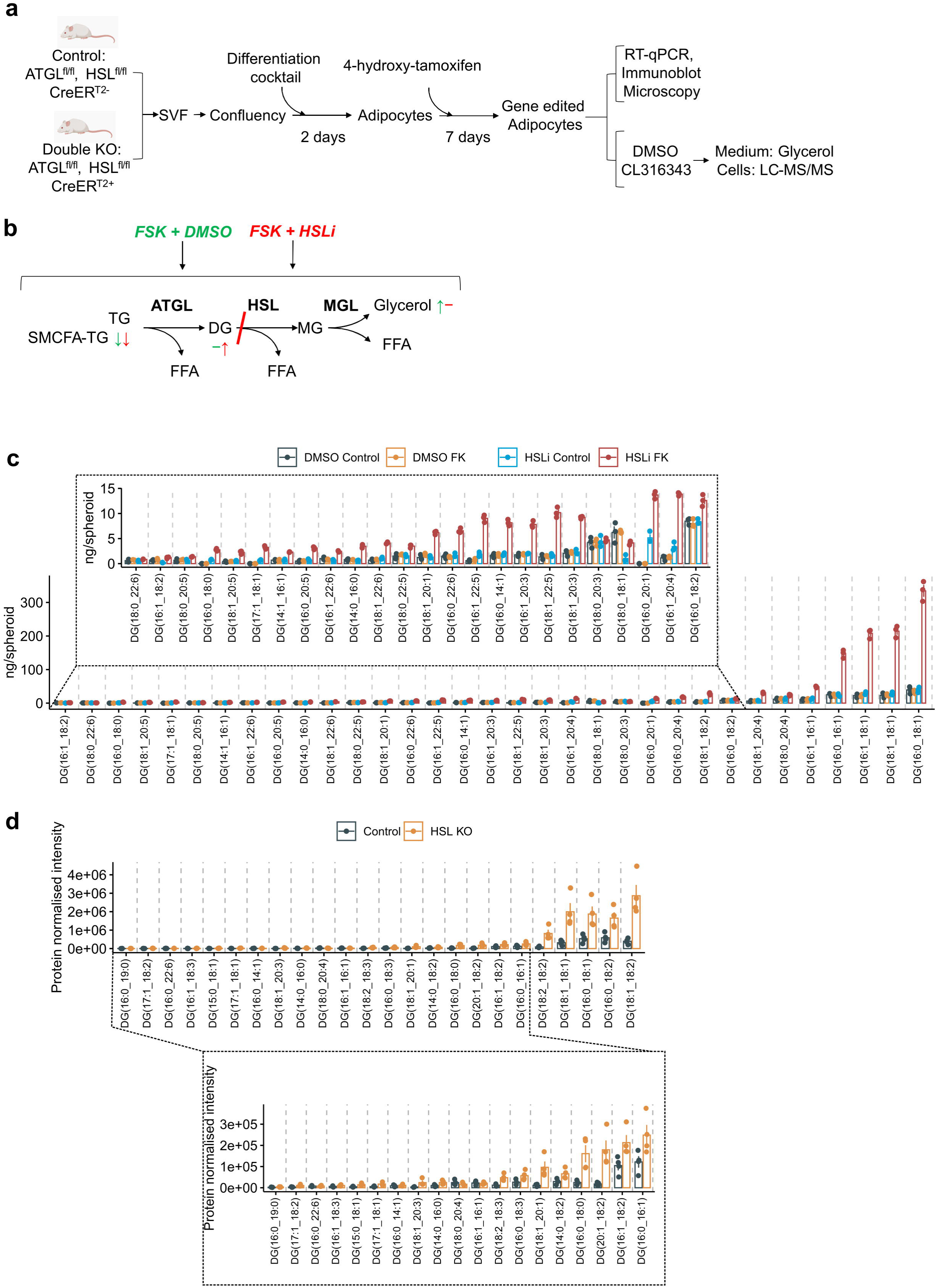
**a**, Schematic overview of the workflow used to prepare lipase knockout (KO) primary adipocytes. **b**, Summary of the stimulated lipolysis experiment performed on DMSO and HSL inhibitor-treated TERT-hWA spheroids shown on **Fig. 5f-j**. **c**, Quantities of each detected DG molecules shown on **Fig. 5j**. Bars indicate mean values of three biological replicates ± SEM. **d**, Quantities of each detected DG molecules shown on **Fig. 5n**. Bars indicate mean values of four biological replicates ± SEM.

